# Synonymous *HTT* CAA/CCA-loss variants associated with early Huntington disease onset enhance toxicity beyond somatic instability

**DOI:** 10.64898/2026.09.03.749261

**Authors:** Glen L. Sequiera, Sophia C. Gjervan, Ryan McCallum, Jia Feng, Oguz K. Ozgoren, Sofia Bergh, Jocelyn Bégin, Jessica Levesley, Hailey Findlay Black, Chris Kay, Tanushri Soomarooah, Larissa Arning, Indhu Shree Rajan Babu, A. Nazlı Başak, Jiří Klempíř, Huu Phuc Nguyen, Åsa Petersen, Michael R. Hayden, Mahmoud A. Pouladi

**Affiliations:** Department of Medical Genetics, Centre for Molecular Medicine and Therapeutics, Djavad Mowafaghian Centre for Brain Health, British Columbia Children’s Hospital Research Institute, The University of British Columbia, Vancouver, V5Z 4H4, Canada; Edwin S. H. Leong Centre for Healthy Aging, The University of British Columbia, Vancouver, V5Z 4H4, Canada; Translational Neuroendocrine Research Unit (TNU), Department of Experimental Medical Science, Lund University, Lund, Sweden; Department of Human Genetics, Faculty of Medicine, Ruhr University Bochum, Bochum, 44801, Germany; Department of Medical Genetics, The University of British Columbia, and Children’s & Women’s Hospital, Vancouver, British Columbia, Canada; Suna and İnan Kıraç Foundation, Neurodegeneration Research Laboratory (NDAL), Research Center for Translational Medicine (KUTTAM), Koç University School of Medicine, Istanbul, Türkiye; Dept. of Neurology and Center of Clinical Neuroscience, Charles University in Prague, 1st Faculty of Medicine and General University Hospital in Prague, Katerinska 30, Prague 2, 120 00, Czech Republic; Department of Psychiatry, Skåne University Hospital, Lund, Sweden; Department of Anatomy and Medical Imaging, Centre for Brain Research, Faculty of Medical and Health Science, University of Auckland, Auckland, New Zealand

**Keywords:** Huntington’s disease, *HTT*, genetic modifiers, cis variants, loss-of-interruption, RAN translation

## Abstract

Huntington disease is a fatal neurodegenerative disorder caused by CAG repeat expansion encoding polyglutamine in the *HTT* gene. Recent studies have shown that loss of CAA/CCA interruptions within polyglutamine-coding CAG tracts and adjacent polyproline-coding region are linked to earlier disease onset. It has been hypothesized that somatic repeat instability, influenced by these interrupted CAG tracts, may mediate this effect. Here we demonstrate that *HTT* CAA/CCA-loss variant linked to early disease onset exacerbates mutant HTT toxicity in cellular models through mechanisms independent of somatic repeat instability. The CAA/CCA-loss variant exhibits significantly higher toxicity than canonical *HTT* sequences in both transient expression and stable cell line models. Notably, this enhanced toxicity persists in knockout cells of the mismatch repair gene MSH3 where somatic instability is blunted, with a consistent toxicity hierarchy (CAA/CCA-loss > CCA-loss > CAA-loss > canonical HTT) in both wildtype and MSH3 knockout cells. Furthermore, the CAA/CCA-loss variant generates elevated levels of repeat-associated non-AUG (RAN) translation products. In HEK293-based cellular models, these results suggest that the disease-accelerating effects of *HTT* CAA/CCA-loss variants involve intrinsic properties of the altered sequence context, highlighting the importance of understanding sequence-specific mechanisms in HD pathogenesis beyond polyglutamine length and somatic instability.

## Introduction

Huntington disease (HD) is a fatal neurodegenerative disorder caused by an expanded CAG trinucleotide repeat, which encodes polyglutamines, in the huntingtin (*HTT*) gene ^1^. The length of the uninterrupted CAG repeat is the primary determinant of disease onset timing, with longer repeats leading to earlier manifestation of symptoms ^2,3^. A growing body of evidence from genome-wide association studies (GWAS) has identified several genetic modifiers of HD, including variants in DNA maintenance genes that influence somatic expansion of the CAG repeat ^2,4,5^. Somatic expansion, the lengthening of the CAG repeat over time in affected tissues ^6^, has been proposed as a key driver of HD pathogenesis ^7,8^. However, recent genetic studies have revealed that not only the length but also the sequence context of the CAG repeat can significantly influence disease progression ^2,9–11^, raising the possibility that mechanisms beyond repeat length contribute to the pathogenesis of HD.

Of particular interest are non-canonical, synonymous sequence variants at the *HTT* CAG repeat locus, specifically the CAA/CCA-loss variant (also referred to as CAG-CCG loss-of-interruption) ^2,5,9–11^. This variant lacks the typical CAA interruption near the end of the canonical CAG repeat tract and has loss of the CCA interruption in the CCG tract. Remarkably, individuals carrying this variant experience disease onset approximately 5 to 15 years earlier than those with canonical repeat sequences of the same length, with this effect being particularly pronounced in those with reduced penetrance alleles (36-39 CAG repeats) ^12^. This makes the CAA/CCA-loss variant one of the strongest known genetic modifiers of HD.

How the CAA/CCA-loss variant affects somatic instability is not clear. While a number of studies have shown no detectable increase in somatic instability, using conventional fragment analysis and small-pool PCR methods, in peripheral blood monocytes or post-mortem brain tissues of CAA/CCA-loss persons with HD ^5,10,11^, the critical question of whether the earlier onset associated with CAA/CCA-loss variant is primarily due to increased somatic instability or if other molecular mechanisms could contribute to their enhanced pathogenicity remains to be experimentally addressed.

In this study, we investigate whether the CAA/CCA-loss variant exhibits intrinsic toxicity independent of its effects on somatic instability. We employ cellular models comparing canonical and CAA/CCA-loss *HTT* variants to assess differences in toxicity profiles and deliberately eliminate somatic instability through genetic approaches. Our findings reveal that the CAA/CCA-loss variant exhibits enhanced toxicity compared to the canonical sequence, even in contexts where somatic instability is blunted in the absence of MSH3. MSH3, a member of the MutS homology family of DNA mismatch repair proteins, has been reported as a genetic modifier in HD by promoting CAG expansion ^4,13^. This supports the notion that additional mechanisms beyond CAG repeat expansion contribute to the heightened pathogenicity of these variants in HD.

## Results

### The CAA/CCA-loss variant exhibits enhanced toxicity in a transient expression model

To investigate the potential pathogenic effects of specific synonymous substitutions in the *HTT* gene, we compared canonical and CAA/CCA-loss *HTT* alleles. We utilized HTT586 fragments, a pathophysiologically relevant species generated by caspase-6 cleavage that has been shown to be critical for cellular dysfunction and degeneration in HD cellular and mouse models ^14,15^. We compared canonical constructs containing 21 (wild type, wtHTT) or 82 (mutant, mHTT-canonical) uninterrupted CAG repeats, to a CAA/CCA-loss variant containing 82 uninterrupted CAG repeats (mHTT-CAA/CCA-loss) (Figure 1A).

**Figure 1.**
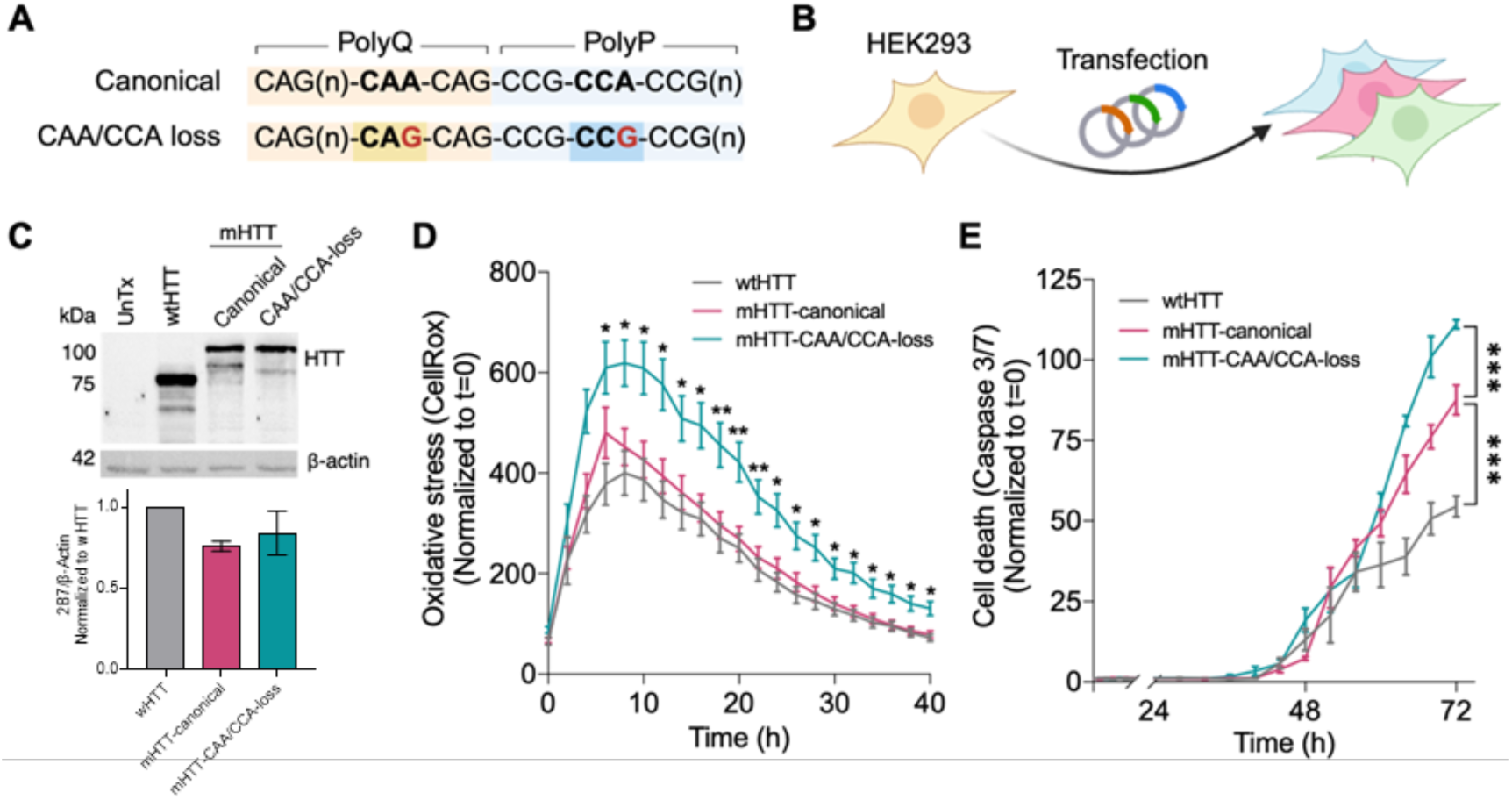
The mHTT-CAA/CCA-loss variant exhibits enhanced toxicity compared to mHTT-canonical in cellular models. (A) Sequence comparison between canonical and the CAA/CCA-loss variant in the *HTT* gene, showing the specific nucleotide differences. (B) Schematic representation of the experimental approach using HEK293 cells transfected with different *HTT* variant constructs. (C) Western blot analysis demonstrating comparable expression levels of HTT protein across untreated WT HEK293T cell line (Untx) wtHTT (wild-type), mHTT-canonical, and mHTT-CAA/CCA-loss variants. (D) Time-course analysis of cellular oxidative stress measured by CellRox fluorescence, showing significantly elevated levels in the mHTT-CAA/CCA-loss variant (teal line) compared to mHTT-canonical (maroon line) and wtHTT (gray line) over 72 hours (n=9). (E) Quantification of apoptosis via Caspase-3/7 activity demonstrating increased cell death in the mHTT-CAA/CCA-loss variant compared to mHTT-canonical and wtHTT over 72 hours (n=3). Error bars represent mean ± SEM; *p<0.05, **p<0.01, ***p<0.001 by two-way ANOVA with Tukey’s post-hoc test.

We established a transient expression system in HEK293 cells to evaluate the relative toxicity of these different repeat motifs (Figure 1B). Western blot analysis confirmed comparable expression levels of the HTT protein across wtHTT, mHTT-canonical, and the mHTT-CAA/CCA-loss variant conditions (Figure 1C). There was no significant difference in HTT levels between the three groups (wtHTT: mean = 1.00 ± 0.00; mHTT-canonical: mean = 0.865 ± 0.170; mHTT-CAA/CCA-loss: mean = 0.915 ± 0.130; one-way ANOVA, F(2,12) = 0.3062, p =0.7419). All experiments were performed as biological replicates (independent experiments on separate days from separate cell preparations).

Cellular stress was assessed using the CellRox fluorescence assay, which revealed significantly elevated oxidative stress in cells expressing the mHTT-CAA/CCA-loss variant compared to mHTT-canonical (Figure 1D). Time-course analysis over 72 hours demonstrated that while both mHTT variants induced greater oxidative stress than the wtHTT control, the CAA/CCA-loss variant produced higher levels of cellular stress (Figure 1D). The difference between the mHTT-canonical and the CAA/CCA variant became apparent within 24 hours of transfection. Consistent with the oxidative stress findings, apoptotic cell death measured by Caspase3/7 activity was also significantly higher in cells expressing the mHTT-CAA/CCA-loss variant compared to the mHTT-canonical (Figure 1E).

Importantly, the enhanced toxicity of the mHTT-CAA/CCA-loss variant was observed within the 72-hour timeframe of transient expression. This relatively short timeline makes it unlikely that somatic instability is the primary driver of the observed phenotypic differences, as significant repeat expansion typically requires longer periods to manifest. Therefore, the increased toxicity of the CAA/CCA-loss variant is more likely attributable to intrinsic properties of the altered repeat sequence rather than primarily a consequence of enhanced repeat instability over this short timeframe.

### The mHTT-CAA/CCA-loss variant exhibits enhanced toxicity independent of somatic instability

To investigate whether the increased toxicity observed with the mHTT-CAA/CCA-loss variant was related to differential somatic instability, we established stable cell lines expressing either the mHTT-canonical or the mHTT-CAA/CCA-loss variant. Using HEK293 Flp-In cells with recombinase-mediated integration, we generated isogenic cell lines with single-copy integration at a defined genomic locus, ensuring comparable expression levels between variants (Figure 2A, Supplementary Figure 1A).

**Figure 2.**
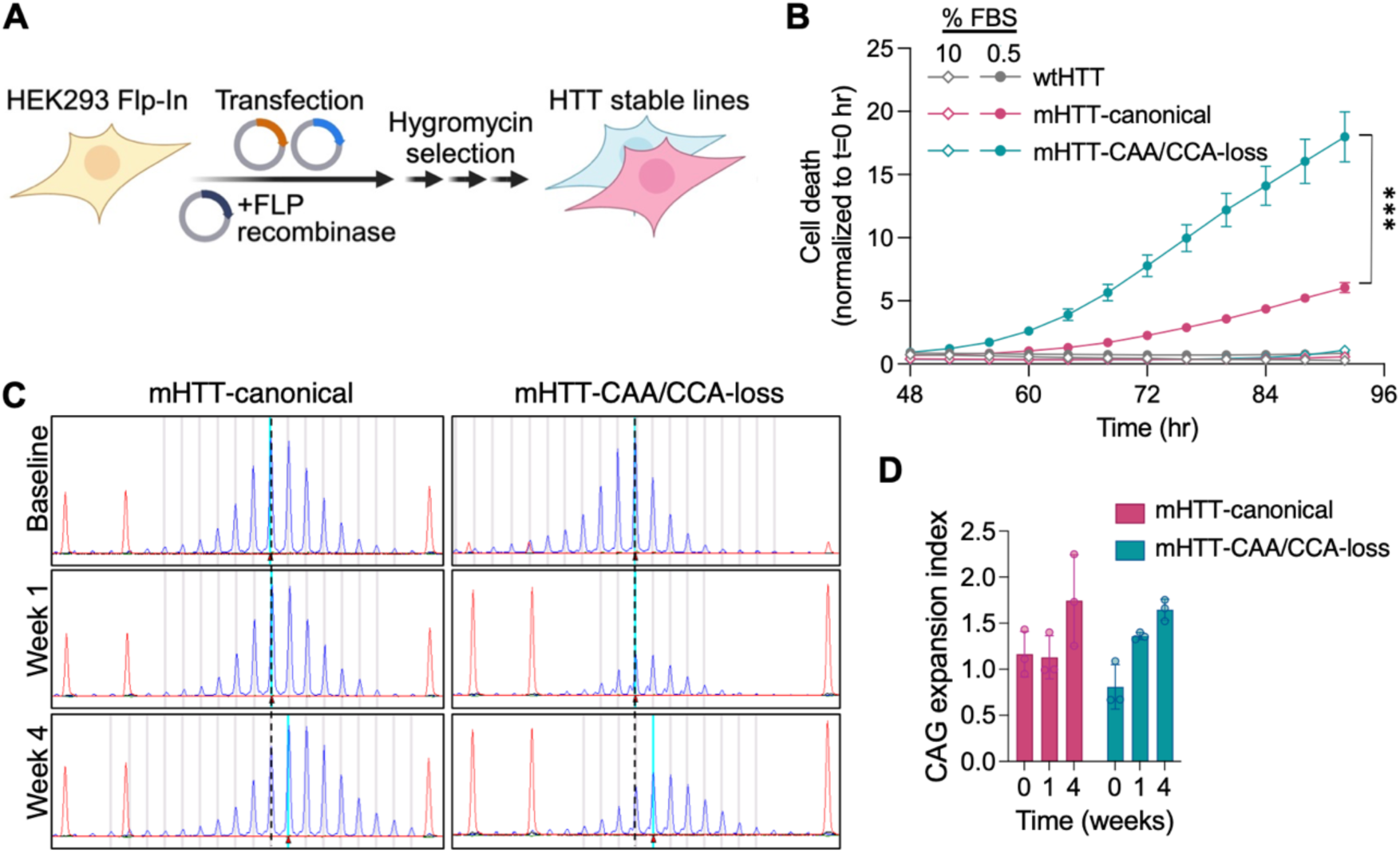
mHTT-CAA/CCA-loss variant exhibits increased toxicity despite comparable somatic instability to mHTT-canonical in stable cell lines. (A) Schematic representation of the experimental approach using HEK293 Flp-In cells with recombinase-mediated integration and hygromycin selection to establish stable HTT expression lines for long-term instability measurements. (B) Significantly increased cell death in the mHTT-CAA/CCA-loss stable line (teal) compared to mHTT-canonical line (maroon) under serum deprivation conditions (0.5% FBS) (n=4). (C) Representative fragment analysis electropherograms tracking CAG repeat stability in mHTT-canonical line (left panels) and mHTT-CAA/CCA-loss line (right panels) at baseline, 1 week, and 4 weeks of continuous culture. (E) Quantification of CAG expansion index over time showing similar patterns of instability between mHTT-canonical and mHTT-CAA/CCA-loss stable lines across 4 weeks, despite the differential toxicity observed in panel B (n=3). Error bars represent mean ± SEM; *p<0.05, **p<0.01, ***p<0.001by two-way ANOVA with Tukey’s post-hoc test.

When subjected to cellular stress through serum deprivation, the mHTT-CAA/CCA-loss stable line exhibited significantly increased cell death compared to the mHTT-canonical line (Figure 2B). Time course analysis revealed a progressive increase in cell death in the mHTT-CAA/CCA-loss expressing cells over 24 hours, while the mHTT-canonical line maintained relatively consistent levels of cell death. Statistical analysis demonstrated significant effects of time (p<0.001), genotype (p<0.05), and a time×genotype interaction (p<0.001), indicating that the mHTT-CAA/CCA-loss variant not only causes higher baseline toxicity but also renders cells more vulnerable to stress over time.

Consistent with the cell death assay, the mHTT-CAA/CCA-loss variant also significantly reduced cell viability as measured by cellular ATP levels in the CellTiter-Glo assay (Figure 2C). Under serum deprivation conditions, the mHTT-CAA/CCA-loss expressing cells showed approximately 40% reduction in viability compared to around 60% viability in cells expressing the mHTT-canonical (p<0.05).

To determine whether differential somatic instability might account for these toxicity differences, we tracked CAG repeat stability in both cell lines over 4 weeks of continuous culture. Fragment analysis revealed similar electropherogram patterns between the mHTT-canonical and the mHTT-CAA/CCA-loss variant at baseline, with comparable changes observed after 1 and 4 weeks of culture (Figure 2D). Quantification of the somatic expansion index showed similar patterns and magnitudes of instability for both variants over the 4-week period (Figure 2E). Both cell lines demonstrated a trend toward increased expansion over time, but no significant differences were observed between the two variants.

These results demonstrate that the mHTT-CAA/CCA-loss variant confers increased cellular toxicity and vulnerability to stress compared to the mHTT-canonical, without detectable differences in somatic instability using fragment analysis over a 4-week period. While the limited sensitivity of these measurements and the relatively short timeframe in HEK293 cells preclude definitive exclusion of instability differences, these results suggest that intrinsic properties of the altered sequence motif contribute to the enhanced pathogenicity.

### Increased toxicity of the mHTT-CAA/CCA-loss variant persists in the absence of MSH3-mediated somatic instability

To definitively determine whether somatic instability plays a role in the enhanced toxicity of the mHTT-CAA/CCA-loss variant, we generated MSH3 knockout cell lines using CRISPR/Cas9 gene editing. MSH3 is a well-established mediator of CAG repeat expansion, and its deletion has been shown to prevent somatic instability in various model systems ^13,16,17^. We first created MSH3 knockout HEK293 Flp-In clonal lines using Cas9 and MSH3-targeted sgRNAs, followed by single cell cloning to isolate homozygous knockout cells (Figure 3A). Western blot analysis confirmed the complete absence of MSH3 protein expression in MSH3KO clonal lines compared to control cells, with -actin serving as a loading control (Figure 3B). The validated MSH3KO HEK293 Flp-In cells were then used as the cellular background for stable integration of either the mHTT-canonical or the mHTT-CAA/CCA-loss variant constructs via FLP recombinase-mediated site-specific integration.

**Figure 3.**
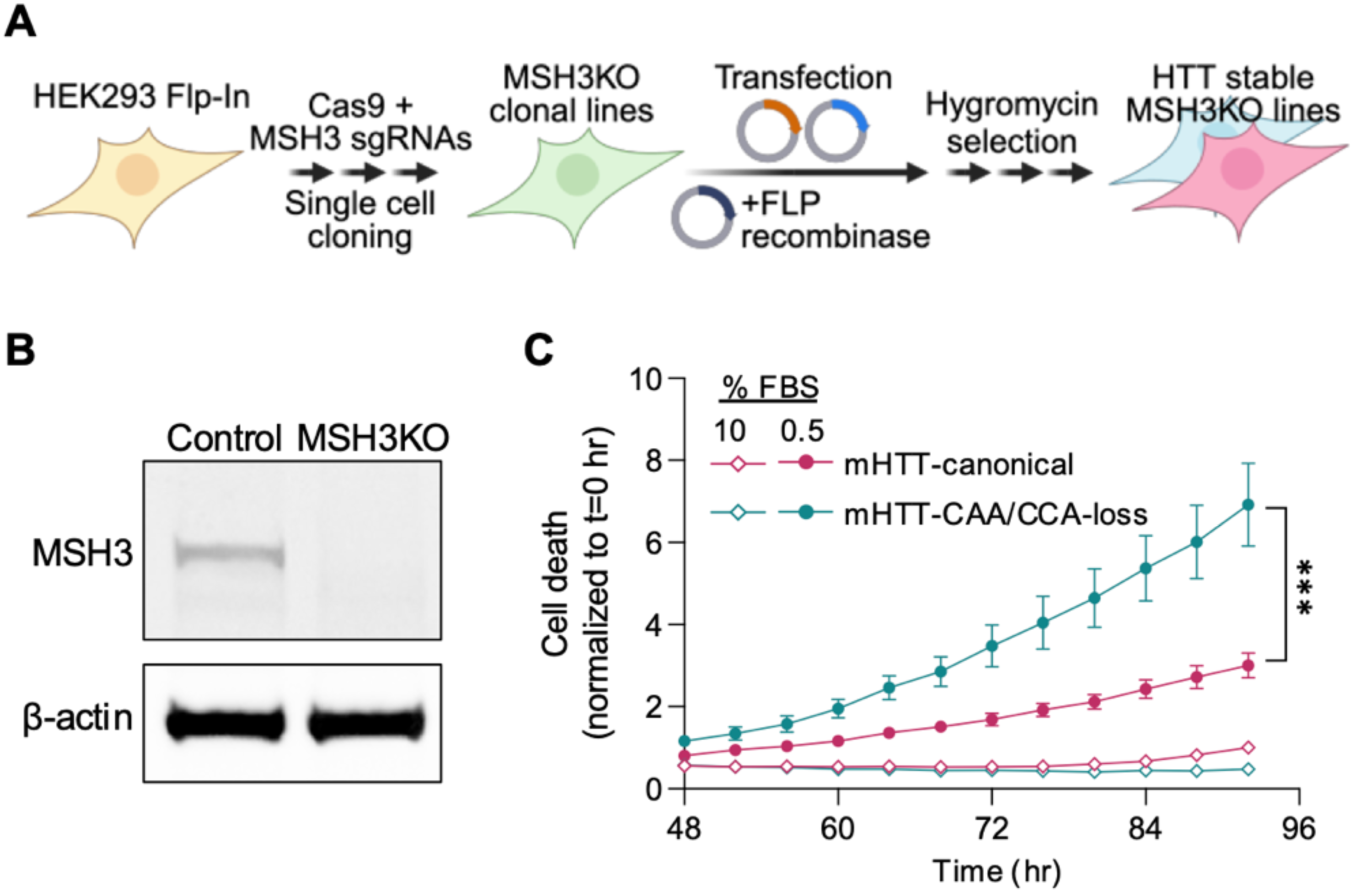
Increased toxicity of mHTT-CAA/CCA-loss persists in the absence of MSH3-mediated somatic instability. (A) Schematic representation of the experimental approach showing the generation of MSH3 knockout HEK293 Flp-In cell lines using CRISPR/Cas9 and sgRNAs targeting MSH3, followed by single cell cloning and subsequent stable integration of *HTT* constructs via FLP recombinase-mediated integration and hygromycin selection. (B) Western blot analysis confirming complete loss of MSH3 protein expression in an MSH3KO HEK293 Flp-In clonal line compared to control cells, with β-actin serving as a loading control. (C) Cell death (NucSpot 680/700 stained area/cell confluency) measured over time (48-96 hr) showing significantly increased cell death in the MSH3KO mHTT-CAA/CCA-loss stable line (teal circles) compared to MSH3KO mHTT-canonical line (maroon circles) following serum deprivation (0.5% FBS, solid symbols) compared to normal serum conditions (10% FBS, open symbols) (n=4). Error bars represent mean ± SEM; *p<0.05, **p<0.01, ***p<0.001by two-way ANOVA with Tukey’s post-hoc test.

Western blot analysis of HTT showed no differences in expression levels of the *HTT* transgenes across the cell lines (Supplementary Figure 1 B). To assess whether MSH3 deletion and the consequent prevention of somatic instability would eliminate the differential toxicity between the two *HTT* variants, we subjected both MSH3KO cell lines to serum deprivation stress. Following serum deprivation (0.5% FBS), the MSH3KO cells expressing the mHTT-CAA/CCA-loss variant showed significantly increased cell death over 48-96 hr compared to MSH3KO cells expressing the mHTT-canonical (Figure 3C). These results provide evidence that the enhanced toxicity associated with the mHTT-CAA/CCA-loss variant occurs independently of MSH3-mediated somatic instability. This result suggests that the CAA/CCA-loss sequence itself confers intrinsic toxicity.

### Evaluating the independent contributions of CAG and CCA-loss variants to HTT toxicity

Other variants besides the CAA/CCA-loss variant have been shown to influence age of onset in HD patients. Both the CAA-loss only variant, which contains a substitution in the penultimate glutamine-coding CAA codon (CAA→CAG), and the CCA-loss only variant, which contains a substitution in the second proline-coding codon (CCA→CCG), are associated with earlier disease onset. The CAA/CCA-loss variant incorporates both substitutions. To dissect the individual contributions of different CAA/CCA-loss variants to HTT toxicity, we designed two additional *HTT* constructs with 82 uninterrupted CAG repeats: mHTT with CAA-loss only or CCA-loss only (Figure 4A). As with previous *HTT* stable lines, these constructs were stably integrated into HEK293 Flp-In cells using site-specific FLP recombinase-mediated integration followed by hygromycin selection (Figure 4B) with similar HTT transgene expression levels across the generated cell lines (Supplementary Figure 1A).

**Figure 4.**
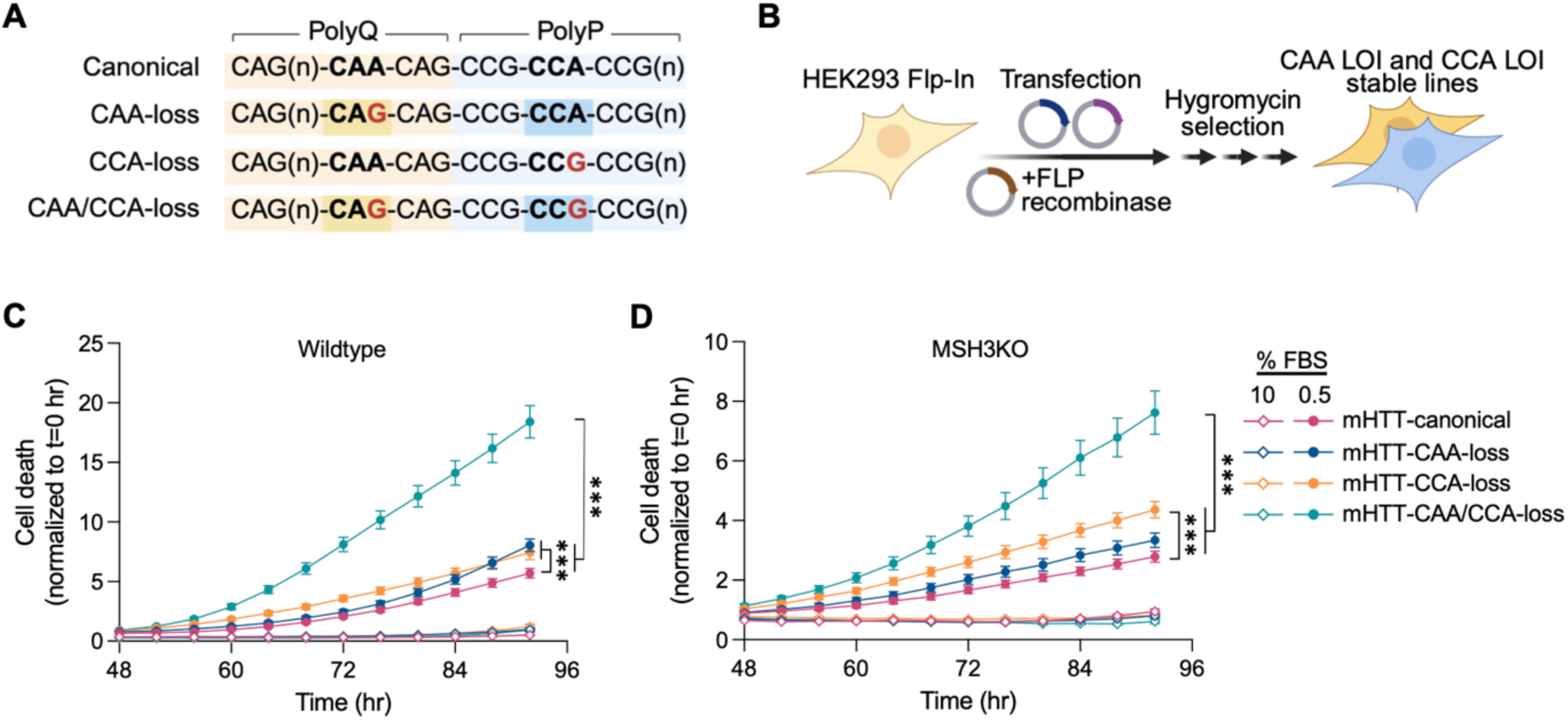
Evaluating the independent contributions of CAG and CCA-loss variants to cytotoxicity. (A) Sequence comparison of canonical HTT repeat structure versus three loss-of-interruption (CAA/CCA-loss) variants: CAA-loss, CCA-loss, and combined CAA/CCA-loss, highlighting nucleotide changes in red. (B) Experimental workflow for generating stable cell lines expressing different *HTT* CAA-loss and CCA-loss variants using HEK293 Flp-In system with hygromycin selection. (C) Cell death kinetics in wildtype cells expressing different mHTT-canonical variants under serum deprivation (0.5% FBS). The CAA/CCA-loss variant (teal) exhibits highest toxicity, followed by CCA-loss (orange), CAA-loss (blue), and mHTT-canonical (maroon) (n=4). (D) Cell death assessment in MSH3 knockout (MSH3KO) cells expressing the same mHTT-canonical variants. While overall toxicity is reduced compared to wildtype cells, the relative toxicity pattern remains consistent: CAA/CCA-loss > CCA-loss > CAA-loss > canonical HTT-82Q (n=4). Error bars represent mean ± SEM; *p<0.05, **p<0.01, ***p<0.001 by two-way ANOVA with Tukey’s post-hoc test.

We first assessed the toxicity of these variants in wildtype cells under cellular stress conditions induced by serum deprivation (0.5% FBS). Cell death measurements over 92 hours revealed a clear hierarchy of toxicity among the *HTT* variants (Figure 4C). The combined CAA/CCA-loss variant exhibited the highest toxicity, reaching approximately 3-fold greater cell death at 92 hours compared to the mHTT-canonical sequence. Individual CAA/CCA-loss variants also displayed enhanced toxicity relative to the canonical sequence, with the CCA-loss variant showing moderately higher toxicity than the CAA-loss variant. Under normal serum conditions (10% FBS), all variants exhibited minimal toxicity, indicating that cellular stress exacerbates the differential toxicity of these variants.

To determine whether these toxicity differences depended on repeat instability, we generated MSH3KO cell lines expressing the same *HTT* variants (Figure 4D, Supplementary Figure 1B). While the overall magnitude of cell death was reduced in MSH3KO cells compared to wildtype cells, the relative pattern of toxicity among the variants remained consistent: mHTT-CAA/CCA-loss > CCA-loss > CAA-loss > canonical. This preservation of the toxicity hierarchy in the absence of MSH3 suggests that the enhanced toxicity of the CAA/CCA-loss variants is not dependent on MSH3-mediated expansion of the repeat tract.

Together, these results indicate that while both CAA and CCA interruption variants independently contribute to HTT toxicity, in combination they produce an additive or synergistic effect. Furthermore, this toxicity pattern persists even in the absence of MSH3-mediated repeat instability, suggesting that the underlying mechanism involves intrinsic properties of the repeat sequence independent of its propensity for expansion.

### The CAA/CCA-loss variant produces elevated levels of toxic RAN translation products

Repeat-associated non-AUG (RAN) translation is a pathogenic mechanism whereby expanded nucleotide repeats drive the synthesis of toxic proteins from alternative reading frames. In HD, aberrant RAN translation has been shown to lead to the production of toxic and aggregation-prone polypeptides, contributing to cellular toxicity and disease pathology ^18–20^. To evaluate whether the enhanced toxicity of the mHTT-CAA/CCA-loss variant is associated with altered RAN translation, we generated stable HEK293 Flp-In cell lines expressing RAN translation reporter constructs (Figure 5A-B), as done previously ^20^. These constructs contained *HTT* exon 1 sequences tagged with epitopes in all three reading frames (RF): Myc for polyQ (RF1), FLAG for polySer (RF2), and HA for polyAla (RF3), enabling simultaneous detection of RAN translation products. Consistent with our previous findings, stable cell lines expressing the mHTT-CAA/CCA-loss variant exhibited significantly increased cell death compared to both wtHTT and mHTT-canonical controls (Figure 5C), confirming that the enhanced toxicity of this variant is maintained in the context of the RAN translation reporter system. Following treatment with 100 nM MG132 (a proteasomal inhibitor) for 60 hours to stabilize RAN products, immunofluorescence analysis revealed that mHTT-CAA/CCA-loss cells had equivalent polyQ (Myc) levels to mHTT-canonical controls (Figure 5D-E). In contrast, the mHTT-CAA/CCA-loss cells showed significantly elevated levels of both polySer (FLAG) and polyAla (HA) RAN translation products compared to mHTT-canonical and wtHTT controls (Figure 5D-E). Automated image analysis quantification demonstrated that the optical density of both anti-FLAG and anti-HA staining, normalized to cell confluency, was significantly higher in mHTT-CAA/CCA-loss expressing cells. These results indicate that the CAA/CCA-loss variant enhanced toxicity that is associated with increased production of RAN translation products in both the polySer and polyAla reading frames.

**Figure 5.**
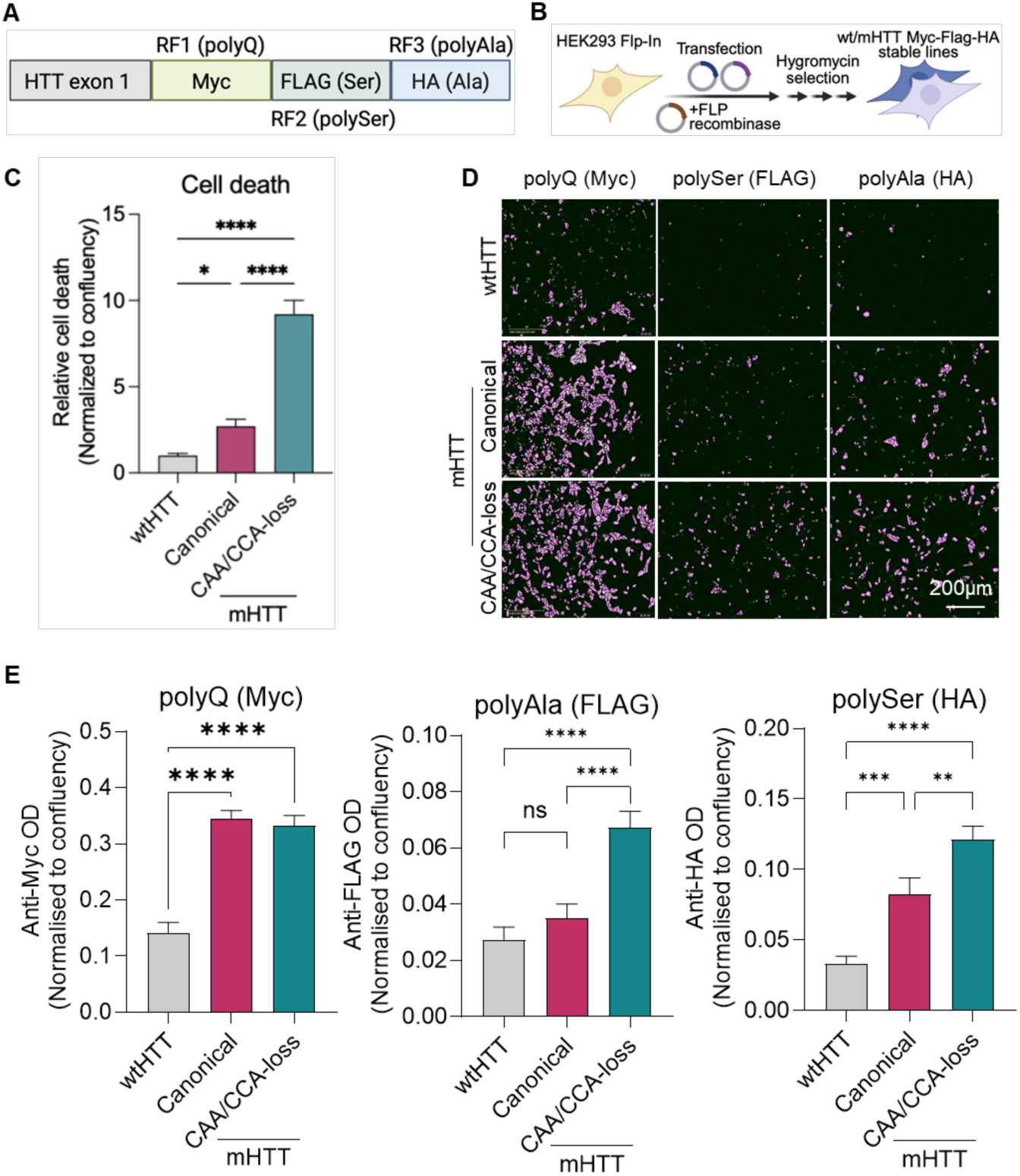
Enhanced toxicity of mHTT-CAA/CCA-loss variant is associated with increased RAN translation. (A) Schematic representation of the RAN translation reporter construct containing *HTT* exon 1 sequence with three reading frames (RF1, RF2, RF3) marked by Myc, FLAG (Ser), and HA (Ala) epitope tags for detection of polyQ, polySer, and polyAla products, respectively. (B) Experimental workflow for generating stable HEK293 Flp-In cell lines expressing wtHTT or mHTT variants with Myc-FLAG-HA tags. (C) Cell death assessment measured by NucSpot 680/700-based toxicity assay in stable lines expressing wtHTT, mHTT-canonical, or mHTT-CAA/CCA-loss variants. The mHTT-CAA/CCA-loss variant exhibits significantly increased cell death compared to both wtHTT and mHTT-canonical. (D) Imaging of RAN translation products in stable cell lines. Representative masked images of positive staining areas for polyQ (Myc), polySer (FLAG) and polyAla (HA) detection in wtHTT, mHTT-canonical, and mHTT-CAA/CCA-loss expressing cells. Scale bar represent 200 μm. (E) Quantification of RAN translation products in stable cell lines. Quantification of anti-Myc, anti-FLAG and anti-HA optical density (OD) normalized to confluency, demonstrating increased RAN translation products in mHTT-CAA/CCA-loss cells compared to controls. Data represent mean ± SEM; ns, not significant; *p<0.05, **p<0.01, ***p<0.001, ****p<0.0001 by one-way ANOVA with Tukey’s post-hoc test.

## Discussion

In this study, we demonstrate that the CAA/CCA-loss variant in the *HTT* gene enhances toxicity through mechanisms that extend beyond increased somatic instability. These findings provide evidence that the pathogenic effects of CAA/CCA-loss variants involve mechanisms beyond increased somatic instability, complementing the established role of somatic expansion in HD pathogenesis.

The CAA/CCA-loss variant represents one of the most potent genetic modifiers identified in HD, advancing disease onset by approximately 5-15 years compared to canonical repeat sequences of equivalent length ^2,5,9,10,21^. This dramatic effect holds even after controlling for polyglutamine tract length in the HTT protein, raising fundamental questions about alternative pathogenic mechanisms. Our cellular models comparing canonical and CAA/CCA-loss variant *HTT* constructs revealed consistently enhanced toxicity profiles for the CAA/CCA-loss variant, even when somatic instability was experimentally controlled for, providing strong evidence for instability-independent pathogenic mechanisms.

Our observation that CAA/CCA-loss variants do not exhibit increased repeat instability compared to canonical sequences aligns with recent reports ^5,10,11^. While these studies did not evaluate cellular toxicity phenotypes, their analyses showed that the CAA/CCA-loss variant was not associated with increased somatic expansion in blood, post-mortem brain samples, or human cellular models. Our toxicity data, combined with these observations, strongly suggest that the disease-accelerating effect of the CAA/CCA-loss variant involves mechanisms beyond enhanced somatic expansion. We acknowledge that survivorship bias may complicate interpretation of instability measurements: if cells carrying the most expanded repeats in the CAA/CCA-loss line preferentially die (as the toxicity data suggest), these expanded alleles would be lost from the population, potentially masking real differences in instability between variants.

A key contribution of our work is the detailed analysis of individual CAG and CCA-loss variants, which provides new mechanistic insights into HTT toxicity. We observed a clear toxicity hierarchy (mHTT-CAA/CCA-loss > CCA-loss > CAA-loss > canonical), demonstrating that both variants independently contribute to enhanced HTT toxicity, with apparent additive or synergistic effects when combined. Critically, this pattern persisted in MSH3 knockout cells where somatic instability was effectively eliminated, providing strong evidence that these sequence variations modulate toxicity through mechanisms that do not require MSH3-mediated somatic expansion. This represents a significant advancement in understanding how specific sequence contexts modulate HD pathogenesis beyond polyglutamine length.

Several molecular mechanisms may explain the instability-independent toxicity of CAA/CCA-loss variants. First, altered repeat sequences may affect RAN translation, potentially generating toxic peptides from both sense and antisense transcripts ^20^. The CAA/CCA-loss variants could enhance RAN translation efficiency or alter the balance of RAN products, contributing to cellular stress independent of canonical polyglutamine toxicity. This is indeed supported by our findings of increased levels of RF2 and RF3 RAN products in cells expressing mHTT-CAA/CCA-loss compared with mHTT-canonical. Second, the modified CAG/CCG patterns in CAA/CCA-loss variants likely form distinct secondary structures in DNA and RNA that differ from those of canonical repeats. These alternative structures could interfere with transcription, splicing, or RNA-protein interactions Specifically, the CAA-to-CAG substitution creates a longer uninterrupted CAG tract that is expected to form a more stable RNA hairpin structure. Since RNA hairpins at CAG repeats have been implicated in promoting RAN translation initiation, this structural change could directly account for the elevated RAN product levels observed in the CAA/CCA-loss variant. Similarly, the CCA-to-CCG substitution extends the adjacent CCG tract, which may form distinct secondary structures including stable hairpins and potentially G-quadruplexes, further modulating RAN translation efficiency ^22–24^, affecting cellular processes even when repeat length remains stable. Third, sequence variations might alter chromatin structure and transcriptional regulation at the *HTT* locus, aspects previously shown to be influence by disease-associated repeat expansions ^25,26^.

Importantly, our results do not exclude a potential role for differential somatic instability as an additional mechanism contributing to the accelerated onset associated with CAA/CCA-loss variants. It is possible for example that very large but infrequent expansions requiring high-resolution sequencing methods for detection could have been missed with the fragment sizing approach used in this study. Rather, these findings highlight a gap in our understanding of sequence-specific toxicity mechanisms independent of somatic instability that warrant further investigation. Notably, we observed that MSH3 knockout substantially reduced absolute levels of cell death under stress conditions, independently of the variant being expressed. This suggests that MSH3 contributes to cellular toxicity through its broader roles in DNA damage response pathways, beyond its established function in promoting CAG repeat expansion.

While using HEK293 cells in this study provided a simplified, non-neuronal background to introduce and compare mutant *HTT* variants, allowing clear elucidation of sequence-specific cellular toxicity, examining the influence of CAA/CCA-loss variants within a neuronal milieu, where RAN translation efficiency and product stability may differ substantially from the HEK293 context, will be of high relevance in future studies. Furthermore, while we examined multiple CAA/CCA-loss variants, other sequence variations at the *HTT* locus may exhibit different mechanisms of pathogenicity that warrant investigation. The 5-15 year clinical onset shift associated with CAA/CCA-loss variants is a substantial effect. The cellular toxicity differences we observe, while consistent and reproducible, are modest in comparison, suggesting that in vivo multiple mechanisms likely combine to produce the clinical phenotype.

Notwithstanding these limitations, our study reveals that the CAA/CCA-loss variant in *HTT* enhances toxicity through mechanisms beyond somatic instability. The distinct contributions of both CAA-loss and CCA-loss variants to toxicity highlight the importance of sequence context in repeat expansion disorders and suggest that for the subset of HD patients carrying CAA/CCA-loss variants (estimated at 7-10% of HD patients, with considerably higher frequencies in populations of African ancestry), targeting somatic instability alone may need to be complemented by approaches addressing intrinsic sequence-dependent toxicity mechanisms. These findings call for a more nuanced understanding of disease mechanisms in HD and potentially other repeat expansion disorders, where sequence context may play equally important roles in pathogenesis.

## Materials and Methods

Cell culture. HEK293 Flp-In™ T-REx™ 293 Cell Line (HEK, Thermofisher, Cat#R78007) were cultured in HEK Medium [Dulbecco’s modified Eagle medium with high glucose (DMEM-HG, Cat#D5671) supplemented with 10% fetal bovine serum (FBS, Gibco, Cat#A5256701) and 1X Antibiotic-Antimycotic (AA, Gibco, Cat#15240062)]. Transient Transfection of HEK cells. HEK293 Flp-In™ T-REx™ 293 cells were seeded at approximately 1 million cells/well of a 6 well plate for WB and 50,000 cells/well of a 24-well plate for imaging in HEK medium without AA. Transfections were performed using Lipofectamine 2000 (ThermoFisher, Cat#11668027) according to the manufacturer’s protocol. Briefly, 1μg (6-well)/ 0.1ug (24-well) of plasmid DNA (HTT586 canonical and CAA/CCA-loss variants) and P2000 reagent were diluted in Opti-MEM (ThermoFisher, Cat#31985070), mixed with Lipofectamine 2000, incubated for 10-15 minutes at room temperature to form DNA-lipid complexes, and then added to cells. Cells were incubated at 37°C with 5% CO2, and expression was analyzed 72 hours post-transfection. Imaging of the live cells was carried out in Incucyte® SX5 under 20x magnification. CellEvent™ Caspase-3/7 Green ReadyProbes™ Reagent (ThermoFisher, Cat#R37111) for apoptosis detection and CellROX™ Orange Reagent (ThermoFisher, Cat#C10443) for ROS detection were added alongside. Image analyses were carried out using 2024B version of the Incucyte software. Cell death was measured as total Caspase-3/7 stained area/cell confluency per image normalized to 0 hrs. ROS was calculated as total CellROX integrated intensity/cell confluency/image normalized to 0 hrs.

Generation of MSH3 knockout HEK cells. MSH3 knockout cell lines were generated using CRISPR/Cas9 genome editing. CRISPR guide RNAs (gRNAs) targeting MSH3 were designed as follows: G1 (reverse strand): 5’-GGCCGCCCGACGCAGGCTTC-3’; G2 (forward strand): 5’-GAGCCGATTCTTCCAGTCTAC-3’; G3 (reverse strand): 5’-CACCTGGTCGGCTGCACCTGTGG-3’. gRNAs were synthesized by Synthego. HEK293 cells were transfected with ribonucleoprotein complexes assembled from TrueCut Cas9 Protein v2 (ThermoFisher, Cat#A36498) and gRNAs using Lipofectamine CRISPRMax transfection reagent (ThermoFisher, Cat#CMAX00008). Following transfection, cells were cloned by limiting dilution using the Pala Single Cell Dispenser, and individual clones were validated by PCR.

Stable expression of *HTT*586 canonical and CAA/CCA-loss variants. HTT586 mHTT-canonical and CAA/CCA-loss variants along with HTT586-wtHTT inserts were cloned into a pcDNA5 Flp-In expression vector (VectorBuilder, Chicago, USA), sequence confirmed through Nanopore sequencing (Plasmidsaurus, Oregon, USA) and integrated into HEK wildtype or MSH3KO-HEK cells by co-transfection with the pOG44 Flp-Recombinase Expression Vector (Invitrogen, Cat#V600520) at a ratio of 1:9 for a total of 2μg for 2.48 million cells using Lipofectamine™ 3000 Transfection Reagent (Invitrogen, Cat#L3000001). Subsequently, selection was carried out using 2.5mg/ml Hygromycin B (Gibco, Cat#10687010). Emergent clones were expanded and validated through western blot and fragment sizing. Cell death imaging was carried out using a membrane-impermeant nuclear dye-NucSpot680/700 (Biotium, Cat#41035) in Incucyte® SX5 under 20x magnification. Image analysis were carried out using 2024B version of the Incucyte software. Cell death was measured as total NucSpot680/700-stained area/cell confluency per image normalized to 0 hrs.

Fragment sizing and somatic expansion index (SEI) calculation. Genomic DNA were diluted to 50 ng/μL. Primers consisted of a forward (HTT_Fragment-size_F: 5’-AAGCTTGGTAGCCACCATGGCGAC-3’) and a 6-FAM-labeled reverse primer (HTT_Fragment-size_R: 5’-CGGCGGCGGCTGAGGAAGCTG-3, spanning the CAG repeat section of the *HTT* exon 1. Each 50 μL reaction contained: 28 μL nuclease-free water, 10 μL 5x Q solution (Qiagen, Cat#210220), 5 μL 10x PCR buffer, 2 μL dNTP mix (Qiagen, Cat#N2050-10-L), 2 μL forward primer (10 μM), 2 μL 6-FAM-labeled reverse primer (10 μM), 0.25 μL HotStar Taq polymerase (Qiagen, Cat#203203), and 1 μL DNA (50 ng/μL). PCR with the following conditions: initial denaturation at 95°C for 15 minutes; 30 cycles of 94°C for 1 minute, 62°C for 90 seconds, 72°C for 2 minutes; final extension at 72°C for 10 minutes; hold at 4°C. PCR products were analyzed by capillary electrophoresis using an Applied Biosystems 3730xl DNA Analyzer. Fragment sizes were determined with GeneMarker V3.0.1. software.

The somatic expansion index (SEI) was calculated for each sample using a Python script available in the Pouladi Lab GitHub repository (https://github.com/pouladi-lab/repeat-instability-calculator). Peak data files (.FSA format) from Week 1 were analyzed to identify the modal peak, defined as the highest peak within the 365–376 bp range. The modal peak size (bp) and height (relative fluorescence units, RFU) were recorded. A threshold for peak inclusion was set at 20% of the modal peak height and applied consistently across all subsequent time points for each sample. Only peaks exceeding this threshold were included in the analysis. The height of each qualifying peak was expressed as a proportion (Mi) of the total cumulative peak height (H). Each peak was assigned a distance value (di) relative to the modal peak, with d = 0 for the modal peak and incrementally higher values for peaks of increasing fragment size. The SEI was calculated using the formula: SEI = Mi di where M0, M1, M2, … represent the proportions of peaks at distances d0, d1, d2, … from the modal peak. SEI is then subsequently calculated as SEI = M0 × 0 + M1 × 1 + M2 × 2 …

RAN translation analysis. Reporter vectors were generated to evaluate RAN translation products by introducing triple tags at the C-terminal in each of the three reading frames (RF) as follows: RF1-Myc (PolyQ), RF2-FLAG (PolySer) and RF3-HA (PolyAla) in the HTT586 mHTT-canonical and CAA/CCA-loss variant inserts, which had been cloned into a pcDNA5 Flp-In expression vector. Plasmids were constructed using Gibson Assembly, following the NEBuilder HiFi DNA Assembly Master Mix (New England Biolabs, Cat#E2621L) kit. Briefly, assembly fragments were designed with 20-40 bp binding overlaps and generated with a combination of polymerase chain reaction (PCR) HotStarTaq DNA Polymerase (Qiagen, Cat#203203) and synthetic fragments generated by Twist Biosciences (San Francisco, CA, USA). Plasmids were transformed into MAX Efficiency Stbl2 Competent Cells (Invitrogen, Cat#10268019), isolated using QIAprep Spin Miniprep Kit (Qiagen, Cat#27104), and screened using whole plasmid sequencing through Nanopore sequencing (Plasmidsaurus, Oregon, USA). Stable HEK lines were established as described before. Cells were cultured at baseline conditions (10% FBS) or 100nM MG132 (proteasomal inhibitor) for 60hrs. Thereafter cells were fixed using 4% paraformaldehyde and stained for the respective tags. Imaging of the live cells was carried out in Incucyte® SX5 under 20x magnification. Image analysis was carried out using 2024B version of the Incucyte software.

Antibodies list. For Western blots: Anti-HTT mouse monoclonal antibody 2B7 (1:1000, CHDI-90000830-5, Coriell), Anti-β-actin (13E5, University of British Columbia, Vancouver) rabbit monoclonal antibody (1:2000, Cell Signalling Technology, Cat#4970), anti-MSH3 Polyclonal Antibody (1:1000, Invitrogen, Cat#PA5-29829). For imaging: Anti-FLAG (clone FG4R, IgG2b) mouse monoclonal antibody (1:200, AbLab, University of British Columbia, Vancouver) and Anti-HA (clone HA.C5, IgG3) mouse monoclonal antibody (1:200, AbLab, University of British Columbia, Vancouver).

Statistical analysis. Data are presented as mean ± standard error of the mean (SEM), except where otherwise stated. Statistical analyses were conducted using Prism 10 (GraphPad) with two-way ANOVA followed by Tukey’s post-hoc test. Differences were considered statistically significant when p < 0.05. level. Data were visualized using box and whiskers plots representing the distribution of staining intensities. For RAN translation comparisons involving only two groups (e.g., canonical vs. CAA/CCA-loss), an unpaired two-tailed t-test was used; for comparisons involving three or more groups, two-way ANOVA with Tukey’s post-hoc correction was applied, consistent with all other multi-group comparisons in this study. Throughout this manuscript, n values refer to biological replicates (independent experiments performed on separate days from separate cell preparations), not technical replicates.

## Data Availability

All data generated or analysed during this study are included in this published article and its supplementary information files. Source data for all figures are provided. The somatic expansion index calculation script is available in the Pouladi Lab GitHub repository (https://github.com/pouladi-lab). Additional data are available from the corresponding author upon reasonable request.

## Acknowledgements

This work is part of the EU Joint Programme - Neurodegenerative Disease Research (JPND) delCAA-HD Project (JPND2023-1822-096). It was supported by grants awarded to M.A.P. from the BC Children’s Health Research Institute (IGAP) and the Canadian Institutes for Health Research (grant #191555), MRH from Canadian Institutes for Health Research (grant #FDN-154278), H.P.N. from the German Federal Ministry of Education and Research (BMBF) under grant #01ED2406, A.P. from the Swedish Research Council (grant #2022/01092 and 2023/01707), the Swedish Brain Foundation (#FO2024-0064), the Swedish governmental funding of clinical research (ALF) at Region Skåne and the Knut and Alice Wallenberg Foundation (# 2019.0467), and J.K. from the European Reference Network for Rare Neurological Diseases (Project ID #739510). A.N.B. gratefully acknowledges the use of the services and facilities of Koç University Research Center for Translational Medicine, and extends also her gratitude to Suna and Inan Kıraç Foundation, Koc University and TUBITAK (Grant no: 124N-070) for their generous support of the study. I-S.R-B. is funded by the CIHR Research Excellence, Diversity, and Independence (REDI) Early Career Transition Award (DI2-190730). M.A.P. holds a Scholar Award, and G.S. holds a Post-Doctoral Fellowship Award from Michael Smith Health Research BC. We thank Katherine van Belois and Dr Costanza Ferrari Bardile for technical assistance, and Drs. Jan M. Friedman and Caroline Benn for valuable discussions.

## Author Contributions

Conceptualization: HPN, AP, MRH, MAP; Formal analysis: GLS, JF, LA, MAP; Funding acquisition: ANB, JK, HPN, AP, MAP; Investigation: GLS, SB, SCG, RM, JF, JB, OO, LA, MAP; Methodology: GLS, JF, LA, ISRB, HPN, AP, MAP; Project administration: HPN, AP, MAP; Resources: HPN, AP, MAP; Supervision: HPN, AP, MAP; Writing – original draft: MAP, GLS; Writing – review editing: GLS, MRH, HPN, AP, MAP.

---

**Supplementary Figure 1.**
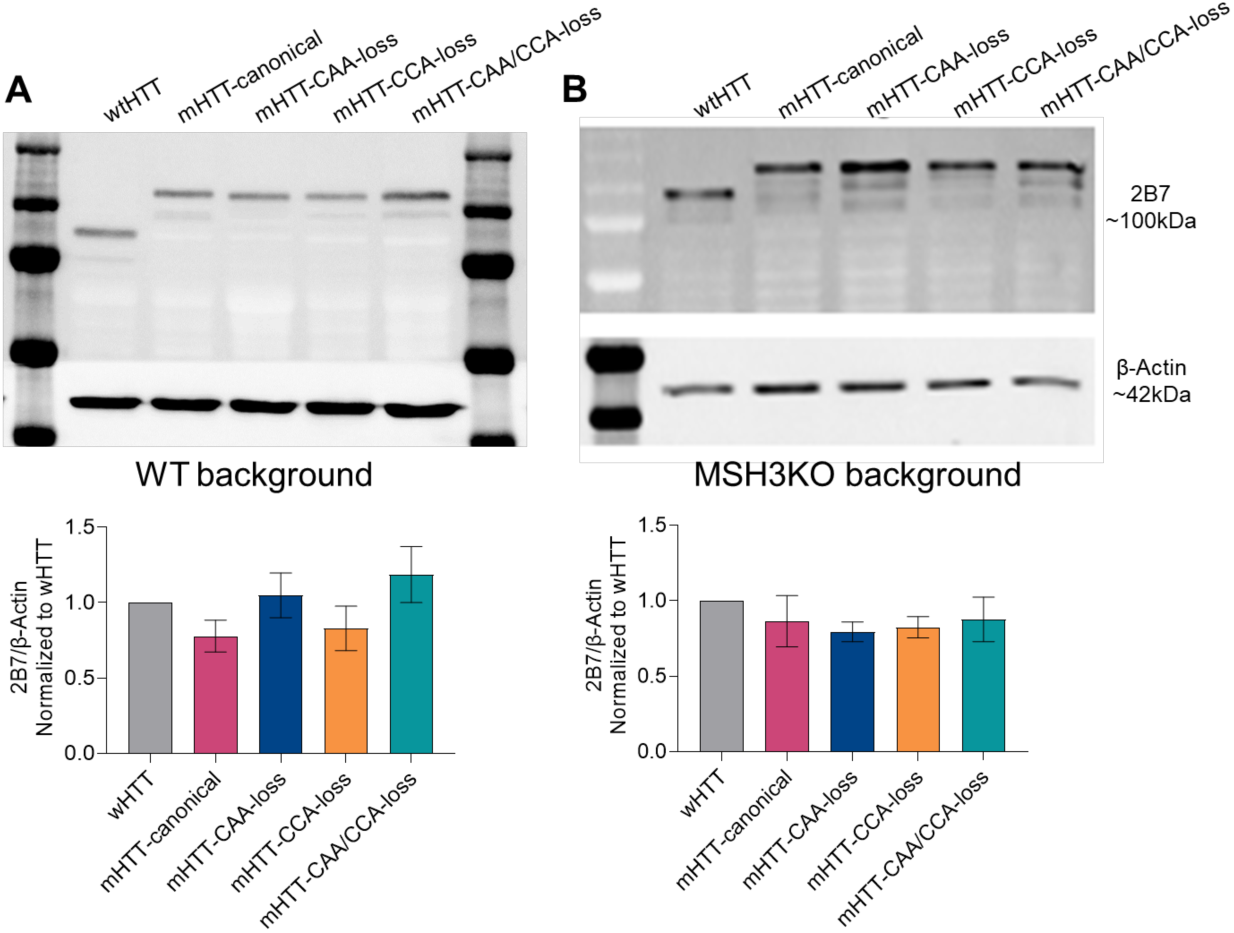
Protein expression analysis of HTT586 constructs in stable HEK293 Flp-In cell lines. (A) Western blot analysis of stable wildtype HEK293 Flp-In cell lines expressing wtHTT, mHTT-canonical, mHTT-CAA-loss, mHTT-CCA-loss and mHTT-CAA/CCA-loss constructs. (B) Western blot analysis of stable MSH3KO HEK293 Flp-In cell lines expressing wtHTT, mHTT-canonical, mHTT-CAA-loss, mHTT-CCA-loss and mHTT-CAA/CCA-loss constructs. HTT protein was detected using the 2B7 antibody. β-actin was used as a loading control. Data represent mean ± SEM, one-way ANOVA with Tukey’s post-hoc test (n=3).

